# Cryo-EM insight into hydrogen positions and water networks in photosystem II

**DOI:** 10.1101/2024.04.02.586245

**Authors:** Rana Hussein, André Graça, Jack Forsman, A. Orkun Aydin, Michael Hall, Julia Gaetcke, Petko Chernev, Petra Wendler, Holger Dobbek, Johannes Messinger, Athina Zouni, Wolfgang P. Schröder

## Abstract

Photosystem II starts the photosynthetic electron transport chain that converts solar energy into chemical energy and thereby sustains life on Earth. It catalyzes two chemical reactions, plastoquinone reduction and water oxidation to molecular oxygen, which both are performed at sequestered sites. While it is known that proton-coupled electron transfer is crucial for these processes, the molecular details have remained speculative due to incomplete structural data. Thus, we collected high-resolution cryo-EM data of photosystem II from *Thermosynechococcus vestitus*. The advanced structure (1.71 Å) reveals several previously unditected occupied water binding sites and more than half of the hydrogen and proton positions of the protein. This unprecedented insight into the structure of photosystem II significantly enhances our understanding of its intricate protein-water-cofactor interactions enabling solar-driven catalysis.

**One sentence summary:** Cryo-EM structure of PSII at 1.71 Å resolution reveals over 50% of hydrogen and proton sites and additional water binding sites, aiding catalytic insight.

## Introduction

By oxidizing water into molecular oxygen, photosystem II (PSII) liberates the electrons and protons utilized in the Calvin-Benson-Bassham cycle for CO_2_ fixation (*1*). Thereby, PSII is at the root of two gas-conversion reactions that have shaped Earth’s bio-, atmo-, hydro- and geosphere (*2, 3*). The unique efficiency of PSII in splitting water with a catalyst made of earth-abundant elements makes it the blueprint for emerging technologies for the sustainable production of fuels and chemicals (*4-6*).

The reaction sequence in PSII starts with the absorption of solar light by the pigments of the associated antenna complexes (Fig. 1). When the excitation energy generated in this process reaches the reaction center of PSII, P680* is formed, which transfers its excited electron to a neighboring pheophytin (Pheo) molecule. Thereby, a strongly oxidizing chlorophyll cation radical (P680^·+^) and a highly reducing pheophytin anion radical (Pheo^·-^) are formed (*7*). This primary charge separation is stabilized by rapid electron transfer steps that increase the distance between the opposite charges and decrease the driving force for their recombination. On the acceptor side, this involves the two plastoquinone (PQ) molecules, Q_A_ and Q_B_. At the water oxidizing (donor) side, P680^·+^ is reduced by the tyrosine side chain Y_Z_, which in turn extracts an electron from a mixed metal oxide cluster comprised of four Mn ions, one Ca ion and five oxo bridges (Mn_4_CaO_5_ cluster), see (*1, 8-10*) for review. By spending about 50% of the chemical energy stored in the P680^*^ (*11*), PSII gains time to wait for subsequent charge separations, generating additional holes and electrons for driving the chemical reactions leading to plastohydroquinone (PQH_2_) formation and water oxidation. Importantly, the Mn_4_CaO_5_ cluster first stores four oxidizing equivalents (S_0_ to S_4_ states) stepwise before templating O_2_ formation from two bound and deprotonated water molecules (*1, 12*).

**Fig. 1.**
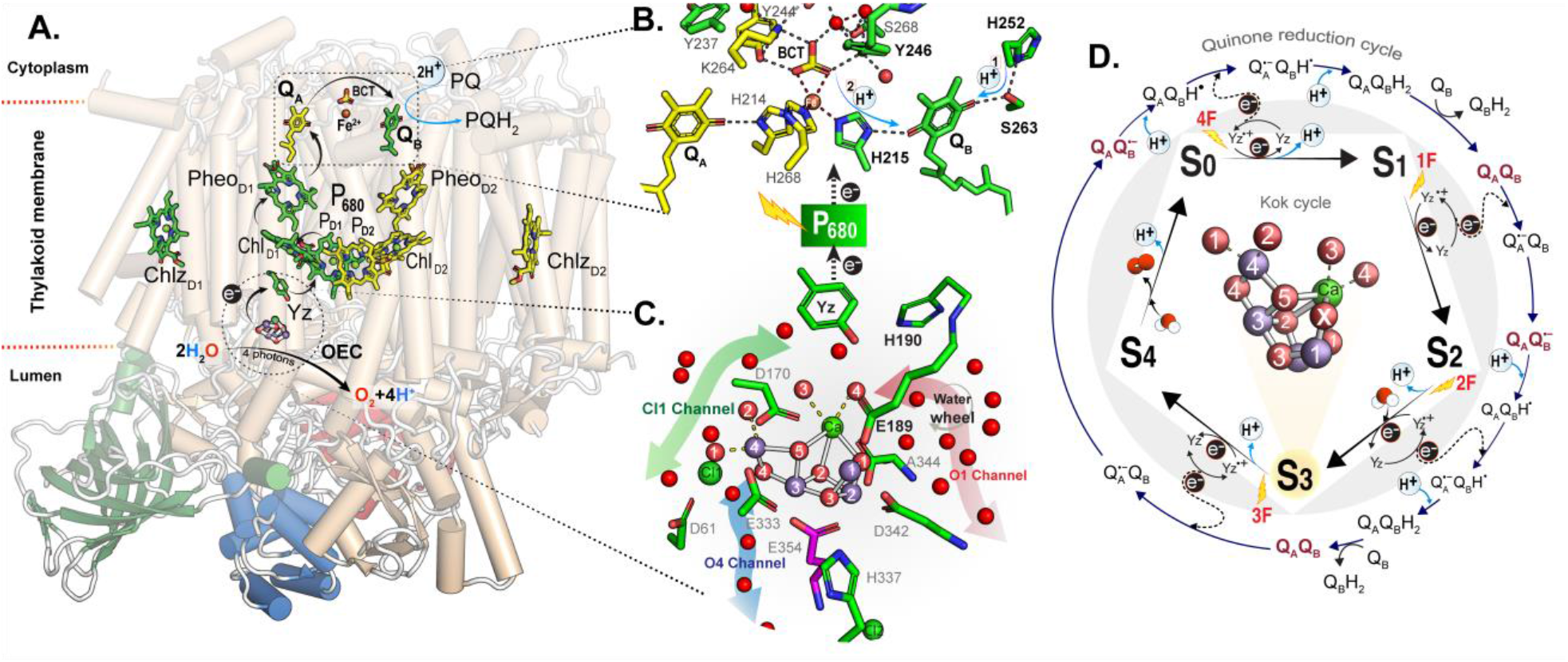
An overview of photosystem II complex and its catalytic centers. **(A)** The PSII structure (one monomer of a dimeric complex is shown) highlights the central redox-active cofactors for charge transfer triggered by light-activated P_680_. The cofactors are colored, while the membrane-embedded helices are shown in light beige. The extrinsic subunits (PsbO, PsbU, and PsbV) are colored green, blue, and red, respectively. **(B)** Detailed view of the acceptor site harboring Q_A_, and Q_B_. A proposed protonation paths of Q_B-_ is indicated (*48*). **(C)** Detailed view of the OEC with its the Mn_4_O_5_Ca cluster and associated water/proton channels (O1, Cl1, and O4) indicated with red, green, and blue arrows. **(D)** Overview of the Kok Cycle for water oxidation at the donor site and its connection to the quinone reduction cycle at the acceptor site. The center shows the Mn_4_O_5_Ca-Ox structure of the S_3_ state (PDB: 7RF8)(*15*). All residues and cofactors in **(A-D)** are colored according to the subunits they belong to: green, yellow, and magenta for D1, D2, and CP43 subunits, respectively. Mn ions, water/oxygen, Ca^+2^/ Cl^-1^, and Fe^+2^ are shown as purple, red, green, and brown spheres, respectively. Black and blue arrows represent electron and proton transfer, respectively.

It is known that proton-coupled electron transfer and local charges are key to the efficiency of the donor and acceptor side reactions of PSII and that water molecules arranged along specific channels play indispensable roles in proton delivery and egress (*10, 13-17*). In addition, internal proton shifts mediated by the proton-water network, such as that between Y_Z_ and the nearby D1-His190, are essential (*18, 19*). Thus, the protonation state of critical amino acid sidechains, as well as the positions and interactions of water molecules, are important pieces to the puzzle of how PSII orchestrates both water oxidation and PQ reduction (*20, 21*).

Recent crystal structures revealed three main channels that connect the Mn_4_CaO_5_ cluster with the lumen: Firstly, the O1-channel, also known as the ‘large’ channel, which starts near the O1 oxo-bridge of the Mn_4_CaO_5_ cluster; secondly, the Cl1 or ‘broad’ channel, which originates from the W1/W2 water ligands of the Mn4 ion and passes along Cl1; lastly, the O4 or ‘narrow’ channel, that begins near the O4 oxo bridge between Mn4 and Mn3 (Fig. 1; for review, see (*16*)). Proton release during the S_2_→S_3_ and S_3_→S_4_ transitions has been shown to occur via the Cl1 channel (*13, 15, 17*), while in the S_0_→S_1_ transition, the O4 channel may be employed (*22-24*).

For the substrate water delivery and binding during the S_2_→S_3_ transition, leading to the formation of an e*x*tra oxo bridge in the S_3_ state that is referred to as Ox or O6, two pathways have been proposed. In one, water (W3) is inserted from the coordination-flexible Ca ion of the Mn_4_CaO_5_ cluster (*25-27*). In this case, fast refilling of the vacant water binding site from a specific part of the O1 channel, known as the ‘water wheel’, has been proposed (*15, 17*). In the other, substrate water delivery through the O4 channel and water insertion via a pivot/carousel mechanism involving the W1/W2 water ligands at Mn4 is suggested (*28-30*).

This emerging picture has been fundamentally challenged by two recent studies. Firstly, on the basis of a reanalysis of published XFEL data, the presence of Ox/O6 in the S_3_ state was challenged (*31*). Secondly, a theoretical study (*32*) concluded that due to crystal pretreatment procedures, none of the present crystal structures presents the physiological hydration state of PSII, making it potentially impossible to draw meaningful conclusions from x-ray diffraction data, especially collected from microcrystals at x-ray free electron lasers (XFELs).

Employing cryo-EM, which involves flash-freezing of fully hydrated PSII samples, we collected high-resolution structural data of dimeric PSII from *Thermosynechococcus (T*.*) vestitus* at a resolution exceeding that of previous cryo-EM and x-ray diffraction studies. Our study confirms the water positions from x-ray crystallography and adds several transiently or partially occupied water positions that support previously proposed substrate water and proton pathways at the donor and acceptor sides. Importantly, the data demonstrate a great potential for detecting light elements such as hydrogen atoms, as demonstrated by the identification of over 50% of the anticipated hydrogen positions of the D1 subunit, greatly facilitating the improvement of structural models (*33*).

## Results

### High-resolution cryo-EM structure of PSII

Structural data of PSII from *T. vestitus* BP-1 was collected to a final resolution of 1.71 Å using single-particle cryo-EM (see Methods and Fig. 2A, Fig. S1 and Table S1). Before freezing, the PSII samples were illuminated with one/two flash(es) to enrich them in the S_2_/S_3_ and Q_B-_/Q_B_ states (see Methods and SI). Both data sets were combined to achieve the best possible resolution.

**Fig. 2.**
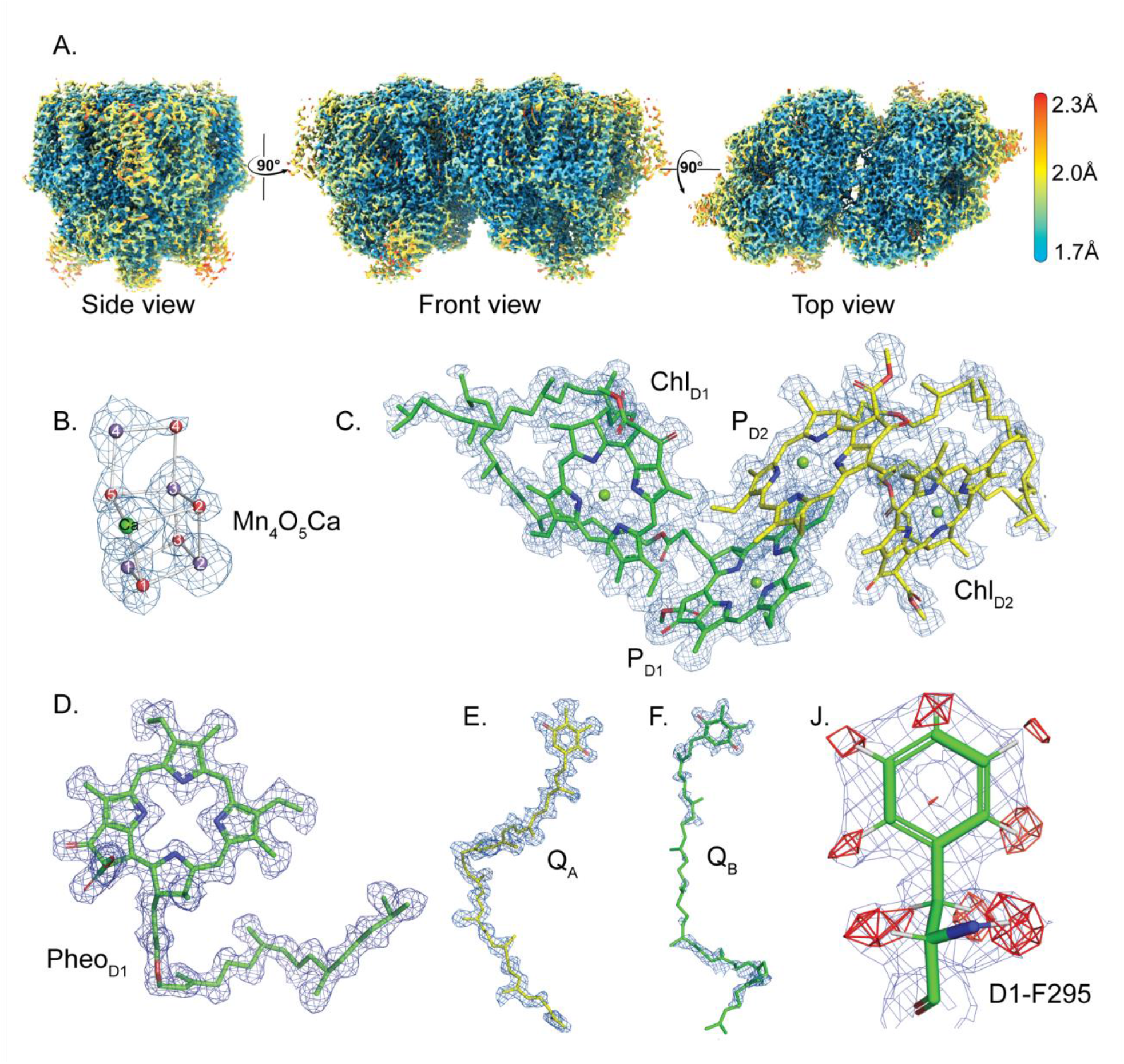
Overall cryo-EM structure of dimeric PSII (PDB …). **(A)** Local resolution map shown at different views of the dimeric PSII shows the resolution quality of the collected data. The map is colored according to the estimated local resolution. (**B** to **F)** show PSII redox-active cofactors with their densities: Mn_4_O_5_Ca cluster, P_680_ (Chl_D1_, P_D1_, P_D2_, Chl_D2_), Pheo_D1_, plastoquinone Q_A_ and Q_B,_ respectively. All densities were shown in blue mesh and contoured at 5 RMSD, except for the heavy metals, which are depicted at 8 RMSD. (**J**) The hydrogen atoms detected in the omitted difference map in one of the amino acid residues of the D1 subunit. The map (shown in blue mesh) and the omit map (shown in red mesh) were contoured at 4 RMSD and 8 σ, respectively.

The refined structure obtained from the combined data set reliably detected the Mn_4_CaO_5_ cluster and other cofactors involved in charge separation and water oxidation (Fig. 2B-F) (*10*). For the Mn_4_CaO_5_ cluster, clear densities were observed for the four Mn ions and the Ca ion, which allows precise modeling of their positions based on the peak maxima. Moreover, all five oxygens connecting the metal ions were directly revealed, removing the need for difference map analysis (Fig. 2B). The structure found is in agreement with previous reports at lower resolution (*10, 23, 34, 35*), but the metal-metal distances are elongated by 0.1-0.2 Å (Fig. S2). This aligns with previous observations that high-valent metal centers are rapidly reduced by electron beam exposure during cryo-EM data collection (*36, 37*) (Fig. S3 and Table. S2). Despite exposing half of the samples to two flashes, no density for Ox/6 was detected (Fig. 2B). We ascribe this to disorder in bridge positions caused by the reduction of the Mn ions (see also O4) and the anticipated low S_3_ occupancy in the merged data set. Nevertheless, the radiation damage in our structure appears to be relatively moderate. Clear densities are evident for all ligands of the Mn_4_O_5_Ca cluster, including D1-A344 with only one conformation (Fig. S4A). Furthermore, only 50% of the disulfide bond between PsbO-Cys19 and PsbO-Cys44 is disrupted (Fig. S4B), unlike in earlier reports. (*36, 37*).

For the acceptor side, the data show a complete density for plastoquinone Q_A_, while for Q_B,_ full density was only resolved at a lower RMSD level of the density values of the map, especially for the isoprenoid tail (see Fig. 2E, F and S5). This is most probably due to the high mobility of the Q_B_ between the two data sets (see below) but may also reflect a lower Q_B_ occupancy.

The current high-resolution cryo-EM map of PSII demonstrates significant potential for elucidating the positions of protons and hydrogen atoms within PSII (Fig. 2J). This enhanced capability is due to the higher scattering potential of electron microscopy for lighter elements compared to X-ray crystallography, which markedly improves the contrast for hydrogen atoms (*38*).

### Water networks enabling water-splitting

The present high-resolution cryo-EM structure of PSII confirms the positions of water molecules in the network around the Mn_4_CaO_5_ cluster suggested previously by room temperature XFEL experiments (*15, 17, 39, 40*) (Fig. 3A). Additionally, it enhances our understanding by revealing sites featuring reduced density due to high mobility and/or partial occupancy, thus requiring high resolution and cryogenic conditions for detection.

**Fig. 3.**
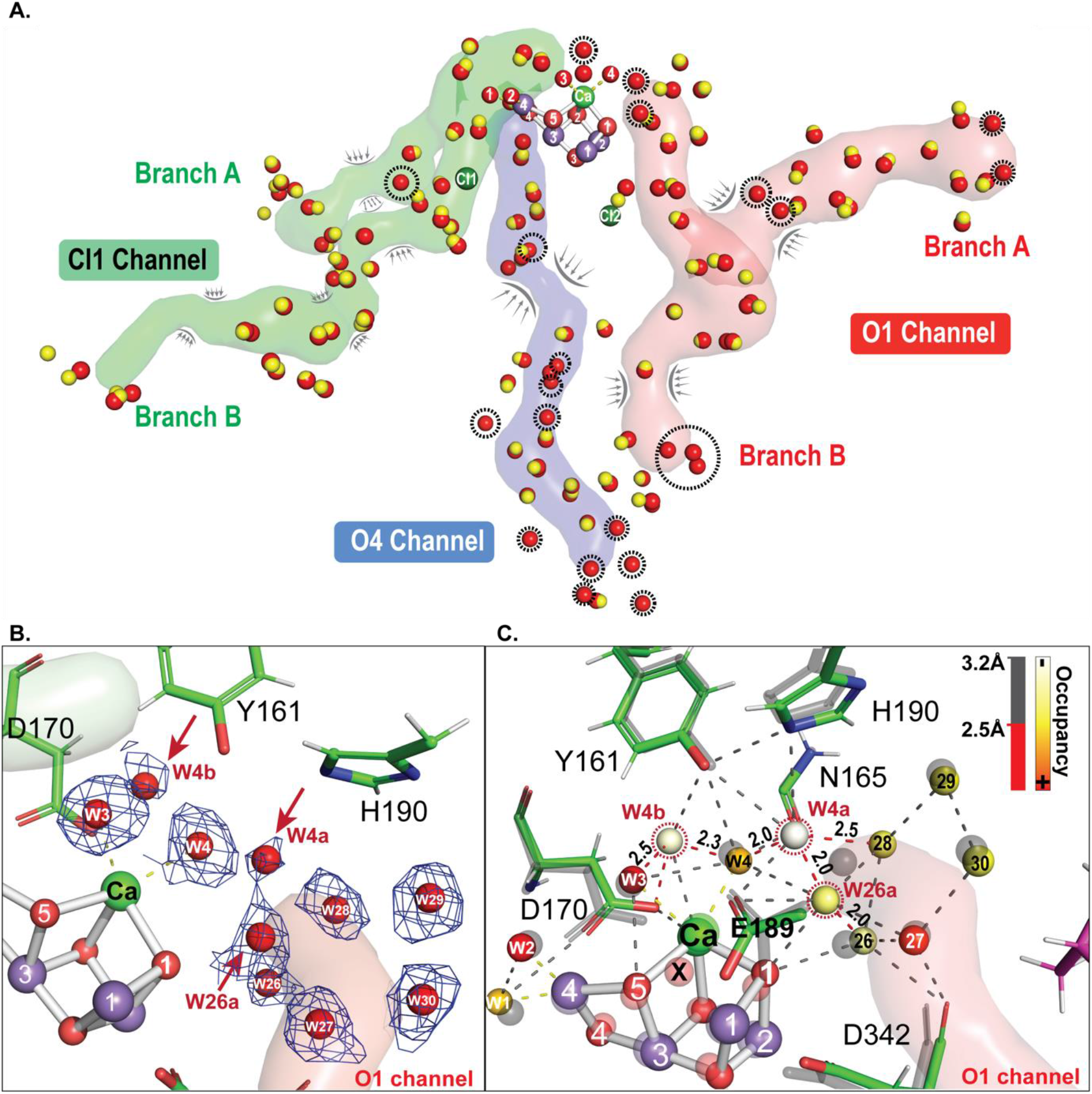
Water molecules in water/proton channels enabling water oxidation by the Mn_4_CaO_5_ cluster. **(A)** The O1, O4, and Cl1 channels are depicted in red, blue, and green, respectively. Water molecules within 5 Å of these channels are shown as red (current cryo-EM structure PDB: …) and yellow spheres (1.89 Å RT-XFEL structure, PDB ID: 7RF1, (*15*)). Waters present only in our present structure are outlined in black dashed circles. Channel bottlenecks are indicated by gray brackets and arrows. **(B, C)** Water molecules in the present cryo-EM structure in the vicinity of the Mn_4_O_5_Ca cluster towards the O1 channel. Red arrows and dotted circles highlight newly detected waters. The density maps of these waters are shown in (**B**) as blue mesh contoured at 3.0 RMSD. In (**C**), the interatomic distances are depicted as dotted, color-coded lines, and the colors of spheres representing water binding sites reflect the occupancy, whereby the lowest value is 0.2 and the highest is 1. The occupancy and the distance scale are shown in the upper right. The RT-XFEL 2F model (PDB ID: 7RF8) (*15*) is displayed in transparent colors.

In comparison to the combined XFEL data set composed of various flashed states (1.89 Å; PDB ID: 7RF1) (*15*), the present data reveal three additional water sites (W26a, W4a and W4b) that are close to the origin of the O1 channel (Fig. 3A-C). Analysis of the hydrogen bond network in this area (Fig. 3C) showed that W26a is hydrogen bonded to O1 and at a very close distance (2.0 Å) to W26, suggesting that it might represent a second position of W26. W4a and W4b are connected to the Mn_4_CaO_5_ cluster *via* H-bonding to W3 and/or W4. We propose that these sites provide more detailed insight into the path of water insertion into the Mn_4_CaO_5_ cluster during the S_2_→S_3_ transition, i.e., from the water wheel to the Ox/O6 binding site located between Ca and Mn1 (*15*) (Fig. 1 and Fig. 3C). We note that a higher positional flexibility of W4 and W3 as compared to W1 and W2 is in line with recent reports (*37, 41, 42*), but we cannot fully exclude structural effects of Mn_4_CaO_5_ cluster reduction.

Going out further in the O1 channel towards the lumen, two additional water binding sites were detected near a narrow section at the entrance to the A-branch. These waters are very close to PsbV-V410 and PsbV-K4; the residues forming the bottleneck of the channel (Fig. 3A). The remaining five additional water sites of the O1 channel are at the exits of the A and B branches, i.e., in areas with very high mobility at room temperature.

In the Cl1 channel, all water molecule positions are consistent with those detected by XFEL (*15*) (Fig. S6). Interestingly, one additional water, W40a, was detected at partial occupancy (Fig. S6B) near the proton gate residues D2-E312, D1-E65, and D1-R334 (Fig. S6B). This water was detectable previously only at the 250 µs time point after the second flash collected by RT-XFEL (*15*). The proximity to W40 and the occupancy pattern suggest that W40a represents an alternative position of W40.

For the O4 channel, 10 additional water molecules were found, but none of those were close to the Mn_4_CaO_5_ cluster (Fig. 3A). The closest one, W55, is around 15 Å away, located just before the first bottleneck region (Fig. 3A and S7A). W55 was detected previously in x-ray diffraction data collected under cryogenic conditions (Fig. S7B and C) (*35, 39, 43*). This indicates high mobility at RT, which may affect the H-network in this area (Fig. S7). Importantly, despite cryogenic conditions and the high resolution achieved in this study, no water molecules were detectable in the subsequent bottleneck region (Fig. 3A). Therefore, our data are not in line with recent MD calculations suggesting that this break is due to artifacts as a result of dehydration of PSII crystals (*32*). Instead, they provide strong support for previous proposals that this bottleneck breaks the H-bonding network of the O4 channel. The remaining 9 additional water molecules were all found past this region, most near the exit of the channel (Fig. 3A). At the start of the O4 channel, W20 is distinctly visible in our data, albeit with a notably lower peak height compared to adjacent water molecules (Fig. S8). This contrasts its absence in previous 1F (S_2_) and 2F (S_3_) XFEL data (*15, 23, 40*), suggesting a significant contribution of S_1_ and S_0_ states to our current structure.

### H-bonding network facilitating plastoquinone reduction

The water-proton network at the acceptor side was investigated using CAVER (*44*). Two main channels, labeled A and B, were detected that connected the non-heme iron (NHI) *via* its ligated bicarbonate (BCT) to the cytosol side of the thylakoid membrane (Fig. 4A). Channel A is comparatively short, wide, and open to the bulk surface through the D1-N247, D2-Q239, and D2-P237 residues (Fig. S9). It harbors several water molecules that form an H-bond network connecting the BCT and the D1-S268 to the bulk (Fig. 4A). Notably, the center path of this wide channel remains largely empty, likely because the waters present there are highly mobile. Interestingly, channel A was previously identified based on MD (*32*) and QM/MM calculations and proposed to provide a path by which BCT/CO_2_ enters to or exits from the NHI, respectively (*21, 45*).

**Fig. 4.**
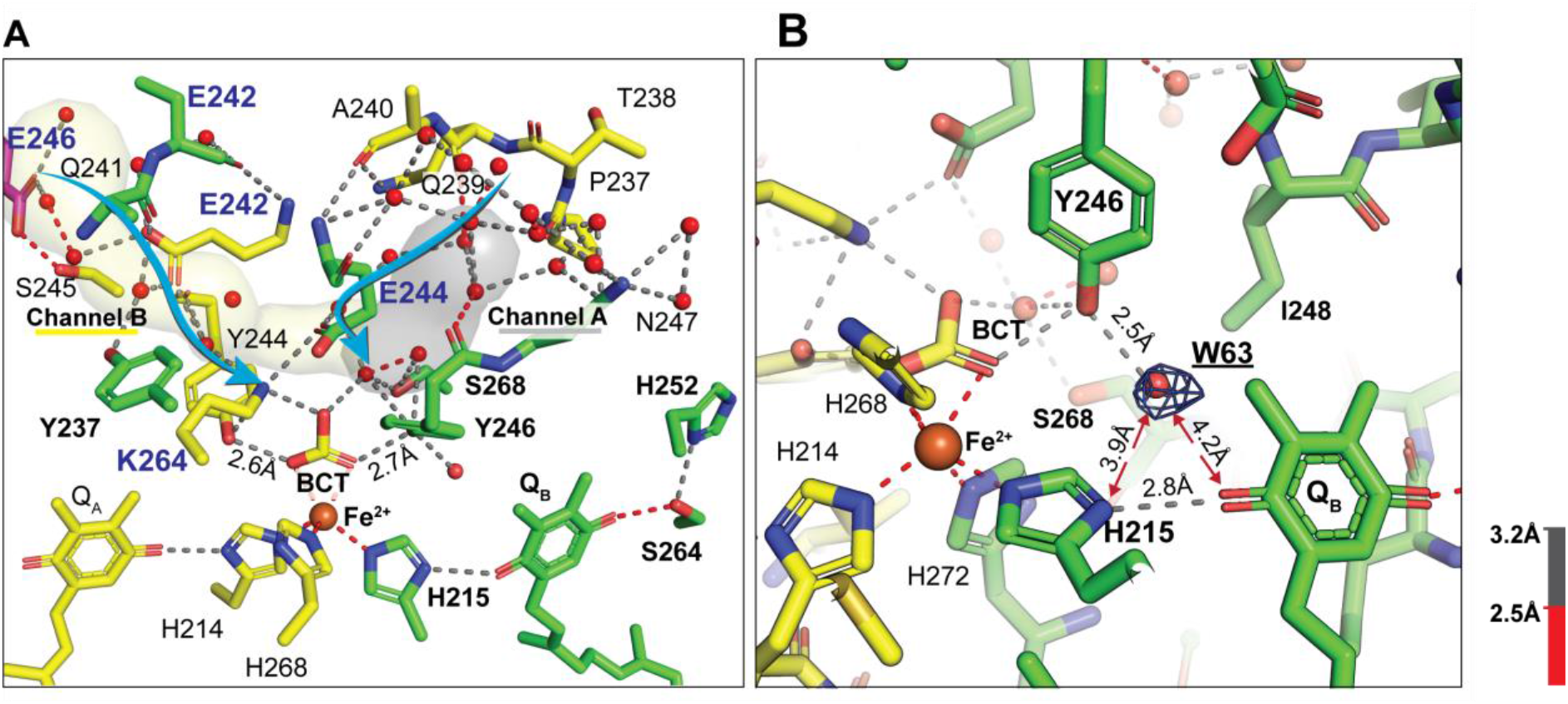
Water/proton channels supporting the acceptor side. **(A)** Water channels are displayed in gray (channel A) and in pale yellow (channel B) were generated by CAVER (*44*) and connect the bicarbonate cofactor (BCT) to the cytosol. The cyan arrows depict suggested protonation pathways of BCT ligated at the non-heme iron. Charged residues were explicitly denoted by being written in dark bold blue. **(B)** Q_B_ binding pocket showing detection of extra density for water W63. The density map of this water is shown as a blue mesh and contoured at 3.0 RMSD. Interatomic distances are color-coded for (**A**) and (**B**), as described at the bottom right. The residues belonging to the D1 and D2 subunits are colored green and yellow, respectively. All water molecules are shown in red spheres.

Channel B is narrow (Fig. S9) and features two bottlenecks that restrict water movements. The first is formed by the BCT, D2-T243, D1-E244, and D2-Y244, while the second is located just before the channel opens to the bulk and is formed by D2-E242, D1-G240, and D2-V247. Channel B comprises several charged amino acid residues and H-bond network analysis (Fig. 4A) suggests that channel B provides a better protonation path for the BCT than channel A. D1-E244, which is a residue found at the branching point of both channels, is proposed to be involved in the protonation of BCT, especially at lower pH (*21, 46*). Recent MD calculations (*32*) report two additional water channels; however, we found no evidence for them, possibly due to their proposed dynamic nature (Fig. S10).

In the Q_B_ pocket, our structure identifies the density for a water molecule, W63, that is situated within a 2.5 Å distance from D1-Y246 (Fig. 4B). Previously, W63 was experimentally observed only for the ΔPsbM mutant of PSII (*47*) and predicted on the basis of QM/MM calculations (*48*), while another theoretical study suggested the presence of two water molecules in this region (*32*). Thus, our finding provides strong experimental support for the proposal that W63 contributes to the re-protonation of D1-H215 *via* BCT and D1-Y246, following the formation of Q_B_H_2_ (*21, 32, 48, 49*).

### Hydrogens and protons

Investigation of the percentage of detectable hydrogen atoms was performed by calculating difference maps with Servalcat (see Methods) (*50*). Using the D1 subunit as a representative example, a value of > 50% was obtained. In the following, some functionally exciting findings are discussed.

Accurate hydrogen positioning refines structural models by reducing side chain rotamer errors, notably for asparagine and glutamine (*33*). For instance, D1-N338, situated at the bottleneck of the O4 channel, has been recently proposed to function as a drawbridge for water molecules in this region based on QM/MM calculations (*41*). Notably, in several previous structures (*19, 23*), the side chain of D1-N338 is reported in a flipped position to the current structure (Fig. S11).

The omitted difference map for protons also identifies a density peak for the shared proton between Y_Z_ and D1-H190 (Fig. 5A). Intriguingly, the peak maximum of this density is closer to the N(ε) of H190 than to the phenolic oxygen of Y_Z_ (1.24 Å and 1.57 Å, respectively), while the total distance of 2.75 Å confirms the presence of a hydrogen bond, albeit weaker than expected (≤ 2.6 Å) (*15, 19*). While the present result clearly illustrates the potential of cryo-EM to address such questions, we note that for a full analysis further studies are required for two reasons. Firstly, in cryo-EM the peak maxima for hydrogen/proton positions are affected by the resolution and B-factor of the dataset and thus typically deviate from their expected locations (Table. S3) (*51-53*). Secondly, the current data may be affected by the reduction of the nearby Mn_4_CaO_5_ cluster (*36*).

**Fig. 5.**
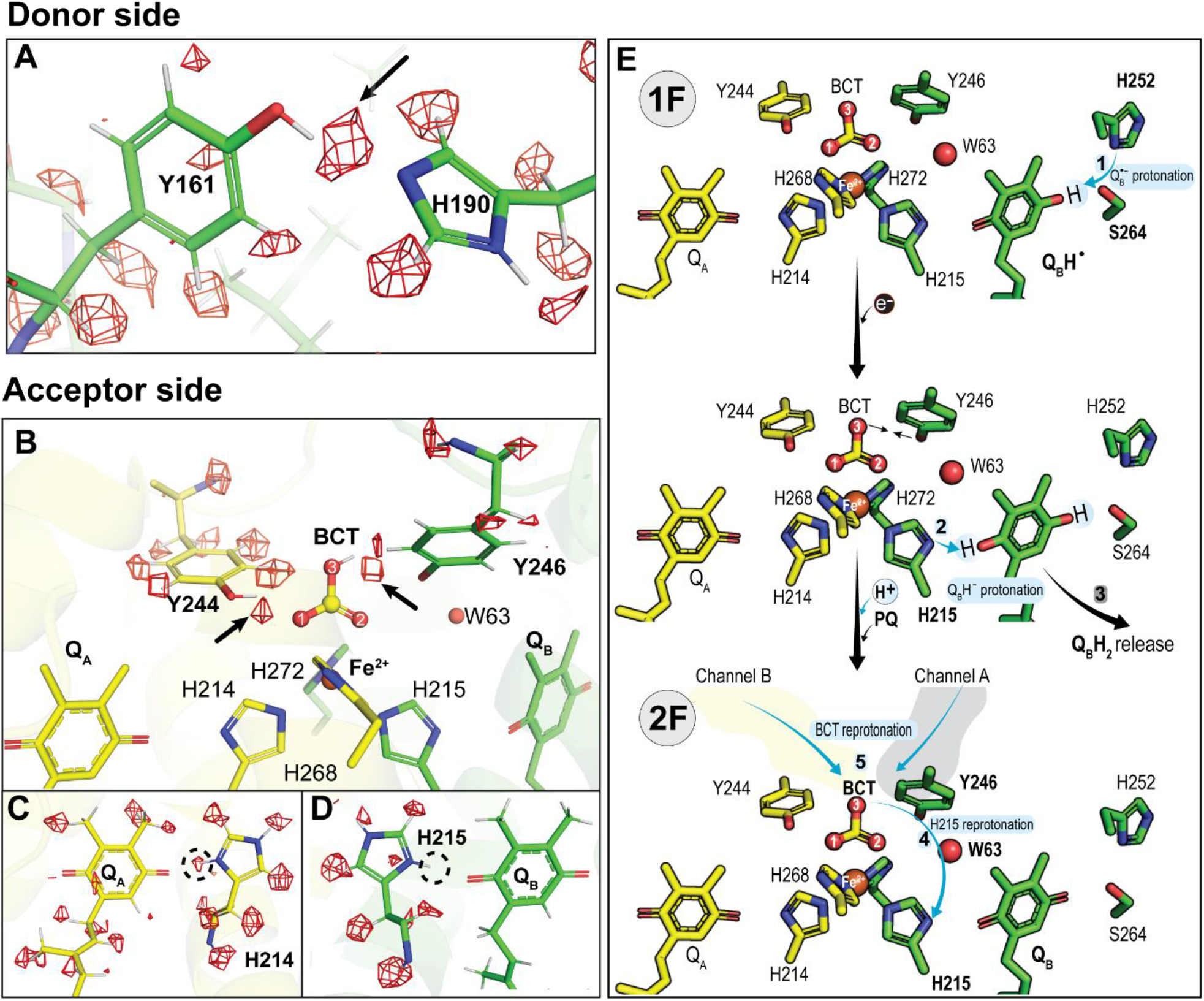
Hydrogens/protons detected at the donor and acceptor sides. (**A-D**) Hydrogen omitted-difference maps of key residues at the donor (**A**) and acceptor (**B-D**) sides. The omit map of the hydrogens and protons are shown as a red mesh and contoured at 8 σ. The black arrows and dotted circles highlight the detected/missed protons at specific residues. (**E**) Schematic summarizing the sequence of events (1-5) leading to the second protonation of Q_B_ that takes place after the second flash. Blue arrows indicate the proton transfer path. The channel A and B connecting the BCT are depicted in gray and pale yellow, respectively.

On the acceptor side, the hydrogen-omitted difference map reveals a density for a proton shared between the phenolic oxygen of D2-Y244 and O1 of BCT, while no density is detected for the proton of the hydroxyl group of D1-Y246 (Fig. 5B). This result corroborates the earlier conclusion that only one of the tyrosine residues, either D2-Y244 or D1-Y246, can form a hydrogen bond with the BCT (*54*). Furthermore, a proton density is also detected near O3 of BCT and in plane with D1-Y246, which has its peak maximum within 1.5 Å from O3 of BCT and 2.3 Å from the phenolic group of D1-Y246. Notably, the distance between O3 of BCT and D1-Y246 is considerably shorter compared to previously reported distances in 0F (S_1_-rich) and 1F (S_2_-rich) structures (*15, 19*) (Fig. S12).

Our analysis showed that all hydrogen atoms and protons of D2-H214, situated close to Q_A_, are detected (Fig. 5C). In contrast, for D1-H215, which is positioned near Q_B_, the proton closest to the proximal oxygen of Q_B_ could not be resolved (Fig. 5D). These findings may indicate that D1-H215 is deprotonated and/or the proton moved to the proximal oxygen of Q_B_ (Fig. 5E). However, the higher B-factor of the Q_B_ region relative to the Q_A_ region (Fig. S12-14), which reflects the differences in the protein environment of the two quinones that enable their disparate functions, results in several undetected hydrogen positions at Q_B_ and its surrounding residues, such as D1-S264 and D1-H252.

Overall, the structural data are consistent with the suggestion indicated (*48*) that Q_B_H^-^ is protonated by D1-H215 and that D1-H215 is re-protonated through D1-Y246 *via* W63 and BCT(Fig. 5E).

## Conclusion

The advancements of cryo-EM have allowed a transformative change in the field of structural biology, enabling us to capture atomic resolution three-dimensional structural data from highly purified and homogeneous protein complexes flash-frozen from their dispersions in aqueous buffer (*55*). While concerns presently remain regarding radiation damage to high-valent metal centers, this should not distract from the exceptional insights provided by this technique.

The present cryo-EM structure (1.71 Å) of dimeric PSII from *T. vestitus* unveils novel partially occupied water-binding sites near the water-splitting Mn_4_CaO_5_ cluster, strongly supporting water delivery to the Mn_4_CaO_5_ cluster *via* the O1 channel. Similarly, it provides the first experimental detection of W63 in intact PSII. By confirming this crucial structural feature of a theoretical calculation (*21, 32, 48*), our data settles the protonation pathways of Q_B_H^-^ and for the subsequent re-protonation of the proton donor (Fig. 5E).

One of the most notable features of cryo-EM is its enhanced scattering potential for light elements, such as hydrogen, which offers distinct advantages over x-ray crystallography at similar resolution. Our analysis shows more than 50% of the hydrogen/proton positions within PSII, which significantly extends present structural models. Knowledge of the protonation states of amino acids, cofactors and/or substrates near the catalytic sites of PSII is essential for revealing how their complex interactions enable efficient proton-coupled electron transfer and water oxidation. Importantly, the various degrees of proton/hydrogen detection in different regions of the protein offer a new path to enhancing our understanding of protein dynamics and their role in catalysis, which is exemplified by the marked differences in the Q_A_ and Q_B_ binding sites.

Therefore, the structural elucidation achieved in this study significantly contributes to the ongoing endeavor of unraveling the molecular basis of the life-sustaining reactions carried out in photosynthesis, thereby refining the blueprints for developing scalable catalysts and devices for producing renewable fuels and chemicals.

## Acknowledgments

The data was collected at the Umeå Core Facility for Electron Microscopy, a node of the Cryo-EM Swedish National Facility, funded by the Knut and Alice Wallenberg, Family Erling Persson and Kempe Foundations, SciLifeLab, Stockholm University and Umeå University. The research was financially supported by Vetenskapsrådet (2020-03809 to J.M.), Germany’s Excellence Strategy coordinated by T.U. Berlin (Project EXC 2008/1-390540038 to A.Z., H.D and P.W.), German Research Foundation (DFG) via the Collaborative Research Center SFB1078 (Humboldt Universität zu Berlin, grant No.TP A5 awarded to A.Z., H.D., R.H.and J.G.) and the Carl Tryggers Foundation (CTS 19.324 to WPS). We acknowledge Drs Mohamed Ibrahim, Asmit Bhowmick, Jan Kern, Junko Yano, and Vittal Yachandra for their helpful discussions.

## Contributions

J.M., A.Z., and WPS designed the experiments, coordinated the research, and provided funding. JG prepared the PSII core complexes under the guidance of A.Z. and A.O.A. P.C developed the flash protocol in the group of JM. A.G. and J.F. prepared the grids with the support of W.P.S.. A.G. and M.H. collected the data. R.H. and A.G. performed the data analysis with the support of P.W. and H.D. R.H. and J.M. wrote the manuscript with input from all authors.

