## Supplementary material for "Cryo-EM insight into hydrogen positions and water networks in photosystem II": Suplematal material

### Materials and Methods

#### Sample preparation

Cultivation of *Thermosynechoccus vestitus* BP-1 was performed as reported previously (*1*). Thylakoid membrane extraction and dimeric PSII core complex (dPSIIcc) purification incorporated minor alterations to the original method. The extracted thylakoid membranes were prepared in a (MCM) buffer containing (20 mM MES/NaOH (2-(N-morpholino) ethane sulfonic acid), pH 6.5, 20 mM $CaCl_2$, 10 mM $MgCl_2$). To get rid of the phycobilisomes, the thylakoid membranes were washed twice using high salt MCM buffer containing (0.3 M $CaCl_2$ and 0.15 M $MgCl_2$). Solubilization was conducted in pH 6.0 at a Chl *a* concentration of 1.7 mM using 0.55 % n-dodecyl-ß-maltoside (βDM) and 0.25 mM Pefabloc® SC-Protease Inhibitor (4-(2-Aminoethyl)-benzyl sulphonyl Fluoride hydrochloride), and for 5 min at room temperature (RT). Purification of the dPSIIcc was then performed using anion exchanger chromatography. Pre-crystallization step was employed to ensure the monodispersity and the homogeneity of the sample. The final purified sample of dPSIIcc was solubilized and stored at 4 mM Chl *a* concentration in a buffer containing (100 mM MES, pH 6.5, 5 mM $CaCl_2$, 5% glycerol, and 0.03% βDM). All procedures were conducted under a dim green light.

The oxygen evolution activity of the purified samples was assessed at RT. The assessment involved measuring the steady-state $O_2$ evolution rate under continuous illumination via a Clarke-type electrode (OxyLab, Hansatech instruments). The dPSIIcc samples exhibited $O_2$ evolution rates of approximately 1,500 μmol $O_2$/(mg Chl *a* h), as measured in buffer containing 50 mM MES/NaOH (pH 6.5), 15mM $MgCl_2$/$CaCl_2$, 1 M Betaine, 2 mM FeCN and, 0.2 mM DCBQ (2,5-dichloro-p-benzoqinone) as artificial electron acceptor at a temperature of 25°C.

#### Vitrification and single particle data collection

For cryo-electron microscopy, 4 µl concentrated (4 mM Chl *a*) dPSIIcc were applied to each glow-discharged (30 s at 15 mA) UltrAuFoil 1.2/1.3 Au 300 grid (Quantifoil Micro Tools GmbH). The sample-on-grid was illuminated with one and two flashes of light (λ = 445 nm, energy pulse = 55 µJ, pulse width = 10 µs, frequency = 2 Hz). Grids were then plunge-frozen in liquid ethane within 194 ± 43 ms from the final flash using an FEI Vitrobot MkIV (Thermo Fisher Scientific) at 100% humidity, 4°C, using a blot force of −5, wait time of 1 second, and blotting time of 5 seconds. All the steps were performed in darkness. Automated data collection was performed using EPU software on a Titan Krios G2 transmission electron microscope (Thermo Fisher Scientific) operated at 300 kV, equipped with a Falcon 4i direct electron detector and a Selectris energy filter at the Umeå Core Facility for Electron Microscopy (UCEM), a node of the SciLifeLab National Cryo-EM facility. From the two grids, a total of 23,324 movies were collected in EER format at a pixel size of 0.57 Å. The total dose was 50 $e^-/Å^{2,}$ and the defocus range was set to -0.8 to -2.0 µm.

#### Single particle analysis

Data collected from the two grids was processed independently but with a similar workflow, using cryoSPARC (v4.0+). In summary, movies were motion-corrected, and CTF-corrected, and resulting micrographs were curated: together, 15,380 micrographs (9,580 + 5,800) with an estimated CTF resolution better than 5 Å, minimal motion (maximum threshold of 20 px) and similar ice thickness were selected for processing. Particles were picked with an elliptical blob picker (minor diameter: 60 Å; and major diameter: 180 Å), extracted with a box size of 720 pixels (Fourier cropped to 180 px), and sorted by 2D classification. Several 2D-classification rounds, followed by ab initio volume generation divided into 3 classes, were needed to achieve a clean particle stack (363,587 + 267,683). Individual particles were motion-corrected, re-extracted at full-box size, and defocus-refined. Non-uniform refinement with CTF-refinement for higher-order aberrations was performed.

The final consensus refinement of each dataset yielded identical 3D reconstructions in the sub-2Å resolution regime. The particle stacks of the independently processed datasets were combined for a joint consensus refinement, with *C2* symmetry imposed, yielding a final 3D reconstruction with an overall resolution of 1.71Å (GSFSC 0.143) (Fig. S1).

#### Post-processing, Model Building, and Structure Refinement

To account for inaccuracies in pixel size, the final electrostatic potential maps were voxel rescaled to best match the scale of a previously published XRD map of a similar sample, PDB 7RF1 (*2*). For the best cross-correlation fit between maps, the pixel size was estimated to be 0.5744 Å. The main map was sharpened with a b-factor of -32.9 $Å^2$. Maps were kept in the same spatial reference, and the PDB model 7RF1 was fitted into the maps using the "Fit in map" function in ChimeraX (*3*). The model-building and refinement processes were conducted using the sharpened map. The structure inspection and editing were conducted using Coot (*4*), and the structure refinement was performed using real_space_refine (*5*) in PHENIX (*6*) with geometric restraints for the protein–cofactor coordination. Metal ions and the oxygens of the $Mn_4O_5Ca$ were placed based on their peak maxima in the sharpened map. The statistics for the data collection, processing, structure refinement and validation are summarized in Table S1. All structural figures were made using Pymol (*7*) or ChimeraX (*3*).

#### Channel analysis

Analysis of water/proton channels at the donor and the acceptor site was conducted by Caver 3.0 (*8*) Pymol plugin. The following settings were used for water/proton channels at the acceptor site: robe radius, 0.9; shell radius, 3.0; shell depth, 4.0; frame weighting coefficient, 1.0; frame clustering threshold, 1.0.

#### Generating the hydrogen-omitted map

The structure was first refined by *REFMAC5* (4) implemented in *Servalcat* (*9*), with refinement settings that permitted the refinement of riding hydrogens and using geometric restraints for the protein–cofactor coordination. Weighted and sharpened $F_o - F_c$ maps were then calculated by *Servalcat* and normalized within the mask, one generated with hydrogens and the other with hydrogens omitted. To enhance the map's clarity for the hydrogen positions, the voxel values below 0.5 were reset to zero for the two weighted $F_o - F_c$ maps. Subsequently, a difference map was then computed between these thresholded maps—one omitting hydrogens and the other including them using the 'make a difference map' function in Coot (3) with a scaling option to normalize the data. The proton and hydrogen analysis was conducted using the generated difference map. The contour mesh levels depicted in all associated figures correspond to 1.172 voxels which is equivalent to 8 σ level within the weighted $F_o$ - $F_c$ map calculated earlier with omitting the hydrogens by the *Servalcat*. The large size of PS II and the estimated presence of approximately 26,124 hydrogens within only one monomer of the protein makes calculating the total hydrogen density detected challenging and exceeding the peak number detection limit of PEAKMAX from the CCP4 suite (*10*). Therefore, The PEAKMAX was used to pick out hydrogen densities ≥7σ exclusively associated with the D1 subunit only as a represented sample for the whole protein. The hydrogen peaks were then filtered by selecting the ones within 1.2 Å from the assigned positions, and in instances where multiple peaks existed at this proximity, the peak with the highest density value was selected. Based on the selected criteria, the analysis revealed that the number of hydrogens linked to the amino acid residue in the D1 subunit is about 1844, representing an estimated 72.5% of the total hydrogen atoms in this subunit (2545), as detailed in Table S3, excluding those hydrogens detected in co-factors.

#### Data availability

The atomic model has been deposited to the Protein Data Bank (PDB) under the accession ID ******** (www.pdb.org). The cryo-EM map has been deposited to the Electron Microscopy Data Bank (EMD) with accession code ****.

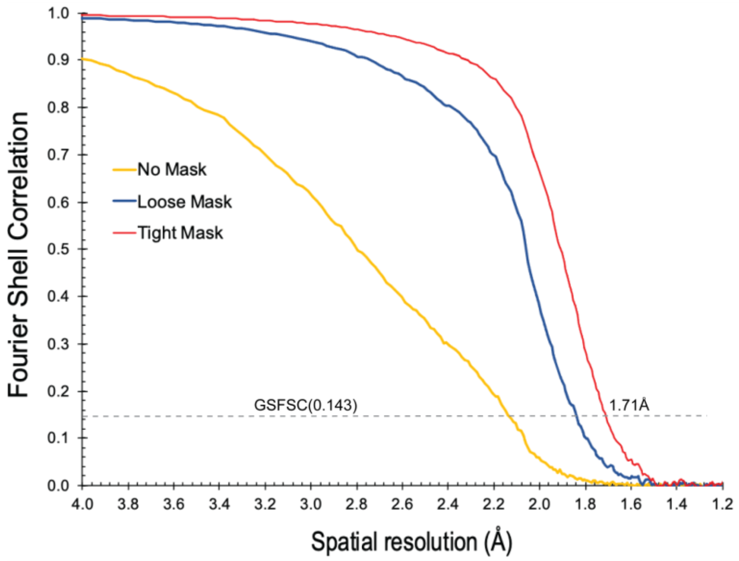

**Fig. S1. Overall resolution estimation of the final map.** Gold standard Fourier shell correlation (GSFSC) curves generated by cryoSPARC at the refinement's last iteration of the final consensus map. The 0.143 threshold resolution is 1.71 Å resolution

**Table S1** Data collection parameters and map/model statistics

| **Data Collection** | |
|---|---|
| Final sample concentration (mM Chl *a*) | 4 |
| Microscope | Titan Krios G2 |
| Detector | Falcon4i |
| Voltage (kV) | 300 |
| Magnification | 215,000 x |
| Pixel Size (Å) | 0.5744 |
| Total electron dose ($e^{-}Å^{-2}$) | 50 |
| Post-column filter | Selectris |
| Defocus range (μm) | -0.8 to -2.0 (in increments of 0.2) |
| Symmertry imposed | C2 |
| Exposure time (s) | 1.9 |
| Dose rate ($e^{-}Å^{-2}s^{-1}$) | 26.32 |
| Dose per frame ($e^{-}Å^{-2}$) | 0.0822 |
| Number of frames per movie | 608 |
| Exposure time per frame (s) | 0.00313 s |
| Number of collected exposures | 13,489 + 9,835 |
| Number of initial particle images | 1,904,999 + 1,121,135 |
| Number of final particle images | 631,270 |
| **Processing and refinement** | PDB (***) |
| Map resolution (Å) | 1.71 |
| FSC threshold | 0.143 |
| Initial model used | 7RF1 |
| Map-sharpening *B* factor ($Å^2$) | -32.9 |
| **Model composition**[a] | |
| Non-hydrogen atoms | 22,796 |
| Ligands | 5544 |
| Waters | 977 |
| **Validation**[a] | |

[a]Sections to be updated based on the final submitted structure.

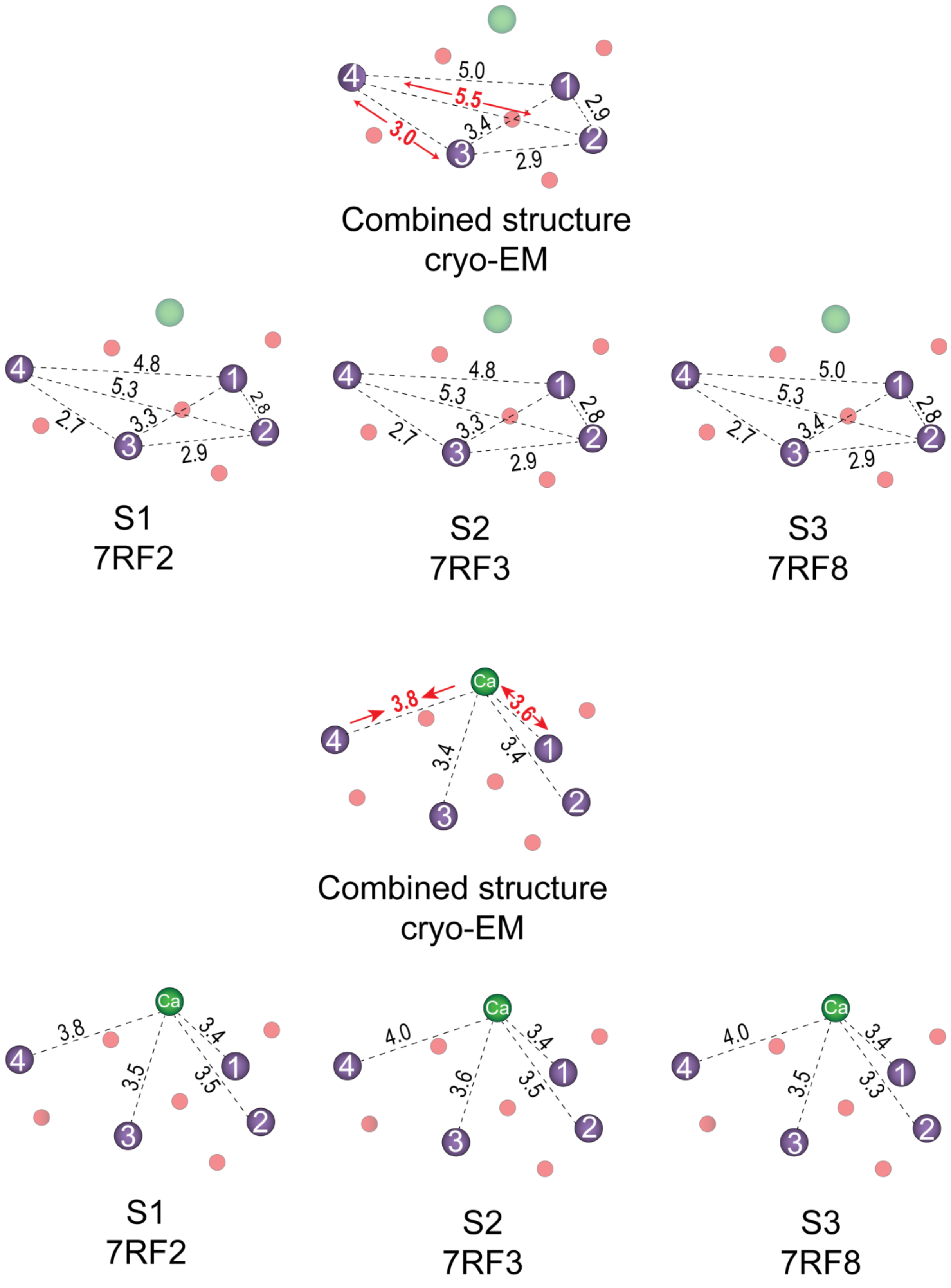

**Fig. S2. The changes in the metal-metal distance (Å) at the OEC of our present combined 1F + 2F cryo-EM data set in comparison to radiation-free structure data collected by RT-XFEL (*2*).**

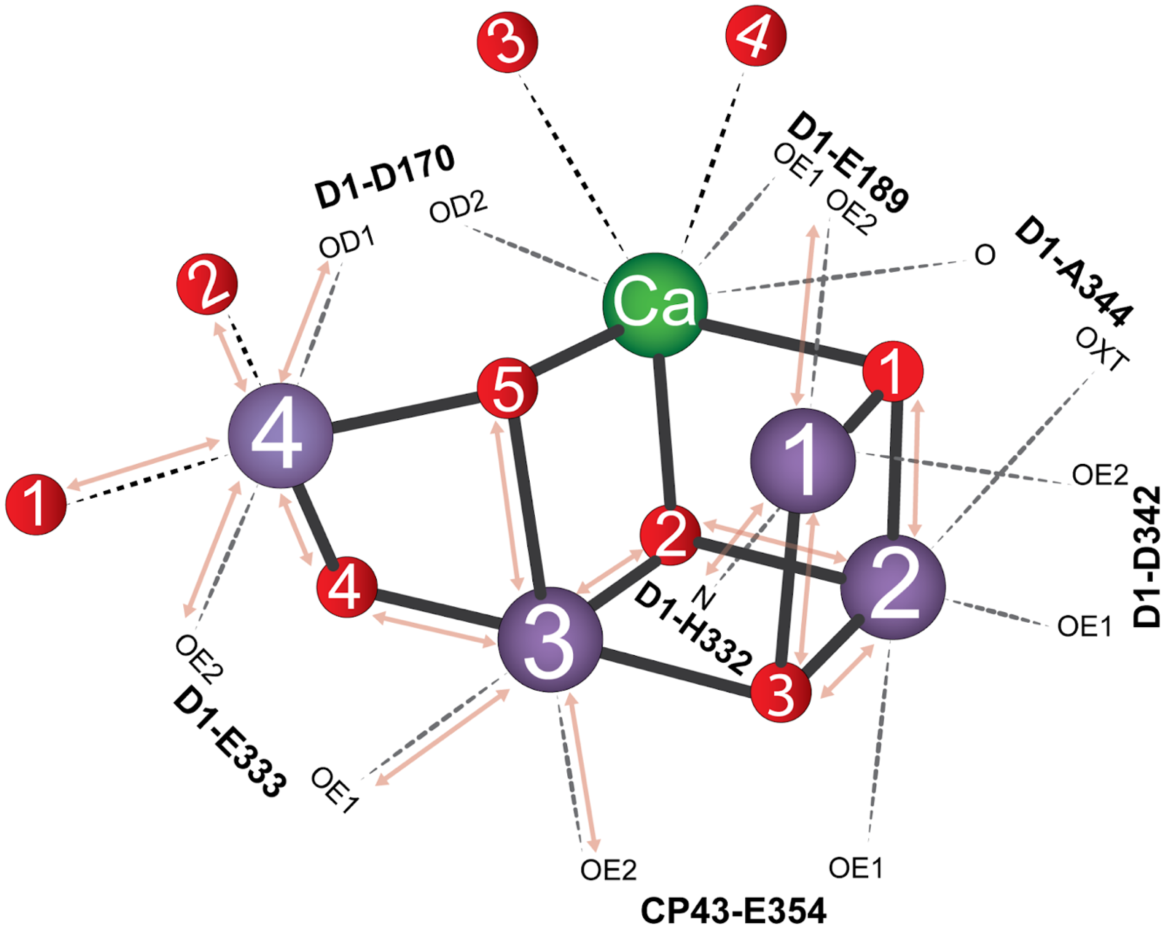

**Fig. S3. Schematic representation of the OEC and its surrounding ligands emphasizing the impact of damage-induced elongation in the distances between metal-ligands in compared to RT XFEL structures (*2, 11*).** The transparent red arrows highlight the major reduction effect detected in Mn1, Mn3, and Mn4.

**Table S2. Distances between Metal-ligands at the OEC in comparison to radiation-free structural data collected by RT-XFEL.** Distances in red represent the distances that significantly increased, by a minimum of 0.17 Å, in comparison to 1F and 2F structure data (*2*).

| | 0F (PDB:7rf2) | 1F (PDB:7rf3) | 2F (PDB:7rf8) | cryo-EM S2-S3 (PDB:… |
|---|---|---|---|---|
| | Distance to Mn1 (Å) | | | |
| O1 | 2.03 | 2 | 1.96 | 2.13 |
| O3 | 1.97 | 1.96 | 1.91 | 2.14 |
| O5 | 2.83 | 2.64 | 2.88 | 2.93 |
| NE2 of H332 | 2.09 | 2.11 | 2.18 | 2.33 |
| OD2 of D342 | 2.16 | 2.05 | 2.01 | 2.17 |
| OE2 of E189 | 1.75 | 1.7 | 1.93 | 2.22 |
| | Distance to Mn2 (Å) | | | |
| O1 | 1.72 | 1.74 | 1.82 | 2.23 |
| O2 | 1.8 | 1.77 | 1.9 | 2.27 |
| O3 | 2.03 | 1.86 | 1.86 | 2.24 |
| OE1 of E354 | 1.84 | 1.97 | 1.84 | 1.9 |
| OXT of A344 | 1.89 | 1.9 | 1.89 | 1.89 |
| OD1 of D342 | 1.97 | 1.96 | 1.98 | 2.1 |
| | Distance to Mn3 (Å) | | | |
| O2 | 1.85 | 1.9 | 1.86 | 2.18 |
| O3 | 1.9 | 1.89 | 2.01 | 2.07 |
| O4 | 1.77 | 1.85 | 1.98 | 2.33 |
| O5 | 1.93 | 1.86 | 1.96 | 2.32 |
| OE2 of E354 | 2 | 1.95 | 2.07 | 2.25 |
| OE1 of E333 | 2 | 1.94 | 1.93 | 2.13 |
| | Distance to Mn4 (Å) | | | |
| O4 | 1.68 | 1.73 | 1.76 | 2.12 |
| O5 | 2.04 | 2.19 | 2.17 | 2.26 |
| W1 | 2.1 | 2.13 | 2.04 | 2.27 |
| W2 | 2.18 | 2.07 | 2.16 | 2.29 |
| OE2 of E333 | 2.06 | 1.78 | 1.96 | 2.18 |
| OD1 of D170 | 1.98 | 1.94 | 1.98 | 2.16 |
| | Distance to Ca (Å) | | | |
| O1 | 2.34 | 2.4 | 2.25 | 2.4 |
| O2 | 2.46 | 2.57 | 2.46 | 2.58 |
| O5 | 2.57 | 2.51 | 2.58 | 2.43 |
| W3 | 2.58 | 2.51 | 2.49 | 2.59 |
| W4 | 2.26 | 2.23 | 2.37 | 2.42 |
| OD2 of D170 | 2.43 | 2.4 | 2.48 | 2.53 |
| O of A344 | 2.59 | 2.52 | 2.55 | 2.47 |
| OE1 of E189 | 2.94 | 2.83 | 3.44 | 3.47 |

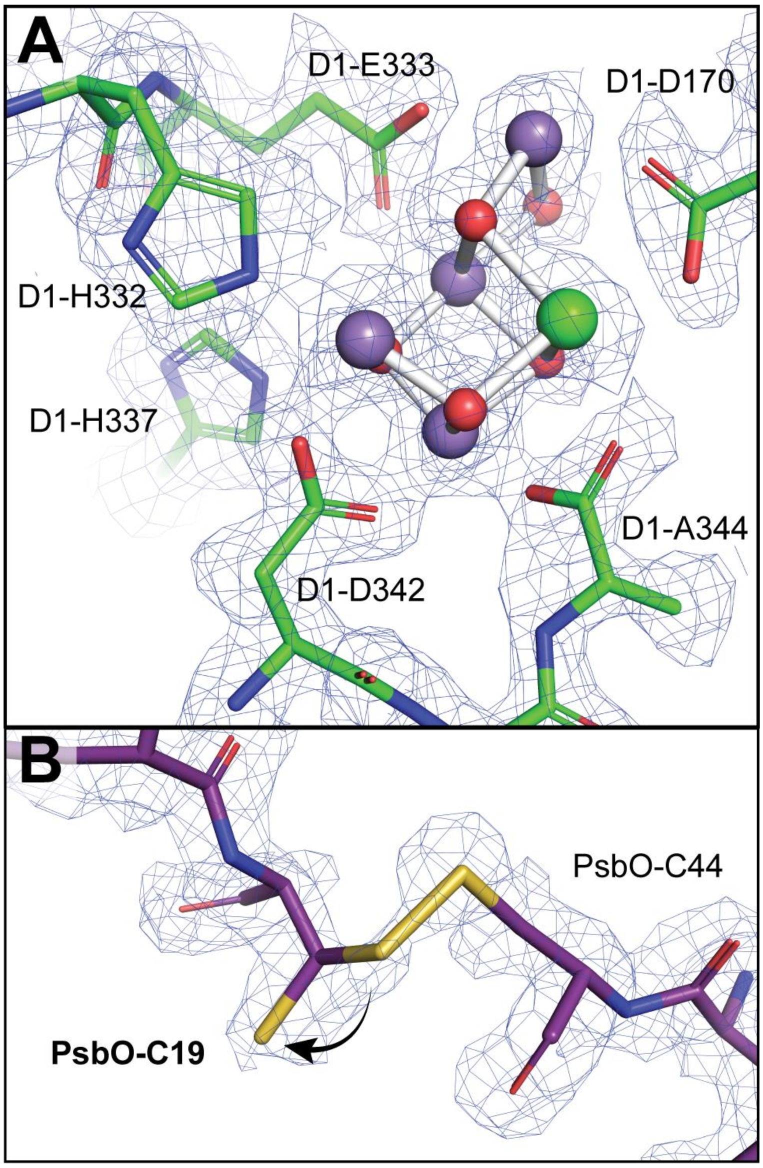

**Fig. S4. (A) The cryoEM Map showing densities of the $Mn_4O_5Ca$ cluster ligands, including D1-A344 in a single conformation. (B) Partial disruption of the disulfide bond between PsbO-Cys19 and PsbO-Cys44.** The black arrow in **(B)** highlights the second conformation of PsbO-C19 side chain with 50% occupancy. The density map in **(A)** and **(B)** is shown as a blue mesh and contoured at 4 RMSD.

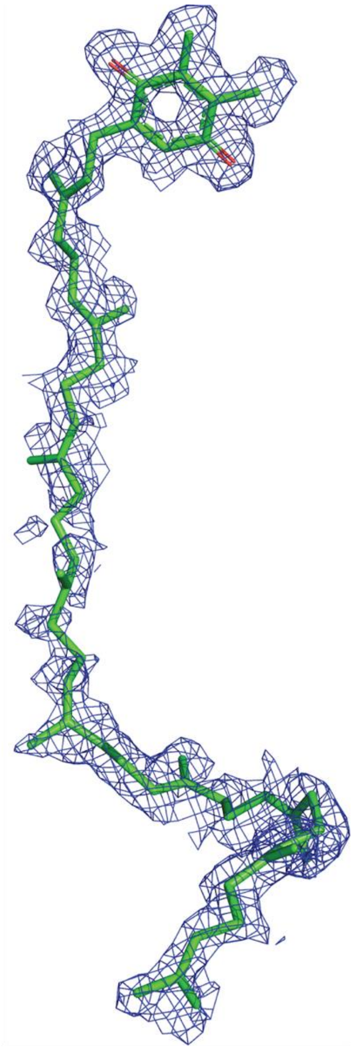

**Fig. S5. The cryo-EM density of plastoquinone $Q_B$.** The $Q_B$ structure with its density contoured at 3 RMSD.

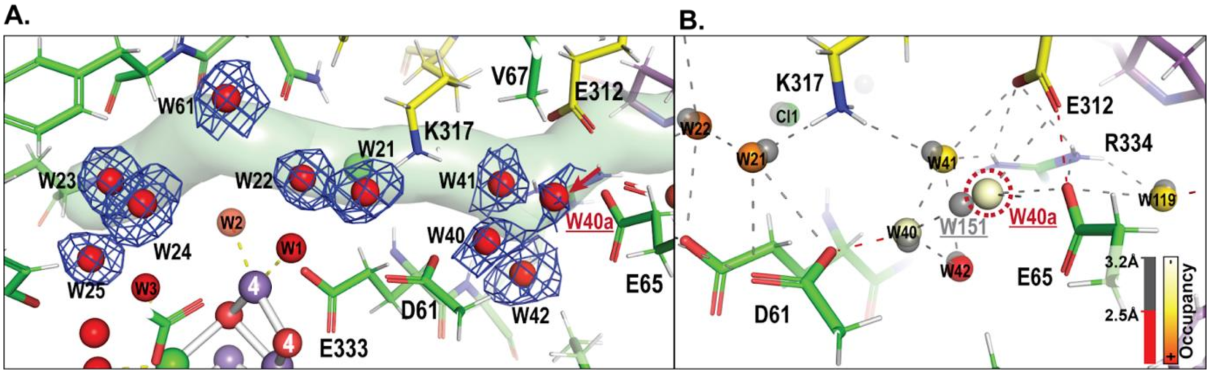

**Fig S6. Water network along Cl1 channel. (A)** Water molecules present in the Cl1 channel. The density map of these waters is shown as a blue mesh and contoured at 3.0 RMSD. The red arrow highlights additional water detected in the current model (PDB ID: …) as compared to the 1.89 Å RT-XFEL structure (PDB:7RF1) (*2*). **(B)** Highlights the H-networks of the additional water molecule detected nearby the proton gate residues E65, E312, and R334. The water molecules are represented using a color gradient scale as described at the bottom right, representing the occupancy of each water, where the lowest value is 0.2, and the highest is 1. The waters from RT-XFEL structure (*2*) (PDB ID: 7RF5, corresponding to 250 µs after the second flash) is overlaid with the model and shown in transparent gray. Interatomic distances are color-coded, as described at the bottom right.

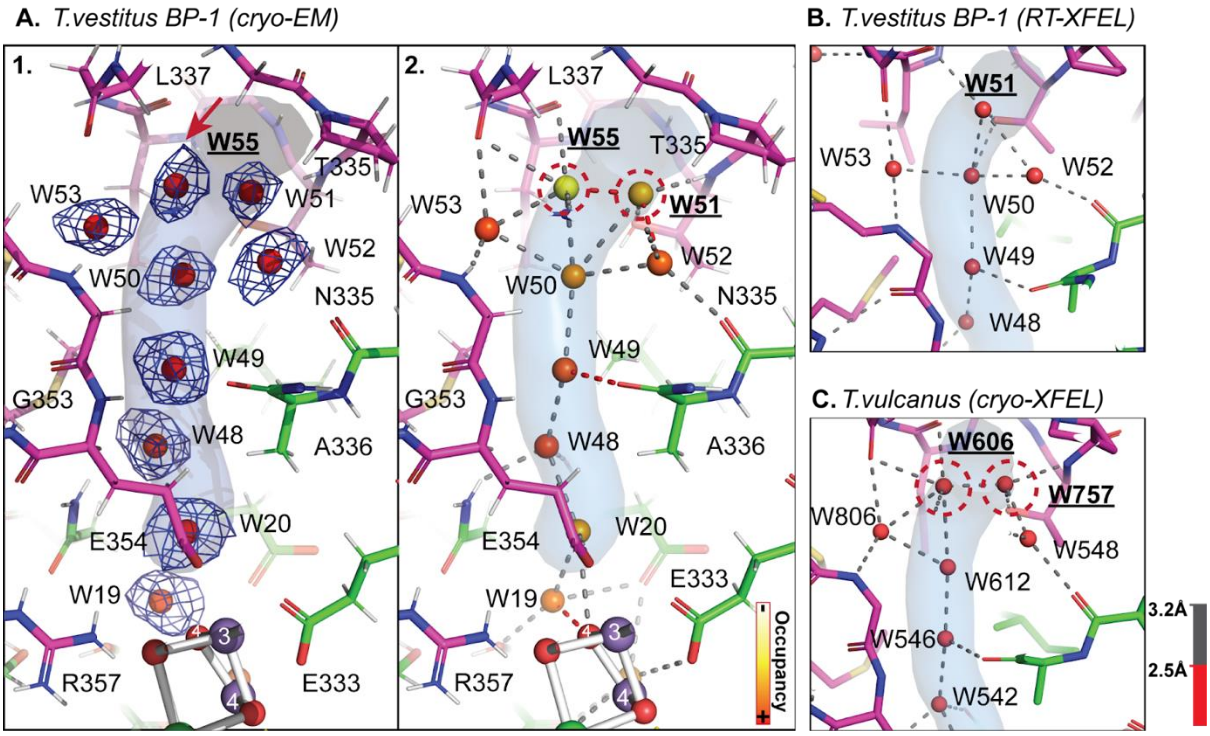

**Fig S7. Water network along the O4 channel. (A)** Water molecules present in the O4 channel. (**A1**) The density map of these waters is shown as a blue mesh and contoured at 3.0 RMSD. The red arrow highlights a water molecule detected in the current model (PDB ID: .. ) that was not identified in RT-XFEL structure measurements (*2, 11*) **(A2)** The H-network along the channel in the current discussed model (PDB ID:..). The red dotted circles highlight the changes in the water structures relative to the RT XFEL structures. The water molecules are color coded, representing the occupancy of each water using a color gradient scale as described at the bottom right, where the lowest value is 0.2 and the highest is 1. **(B)** The H-network along the channel in the RT XFEL structure(*2*) (PDB ID: 7RF1, corresponding to high-resolution RT XFEL1.89 Å structure)). **(C)** The H-network along the channel in the cryo XFEL structure (PDB ID:4UB6 (*12*), corresponding to the dark state rich structure; 0F)). The red dotted circles highlight the difference in the water structure. For (**A**), (**B**) and (**C**), Interatomic distances are color-coded, as described at the bottom right.

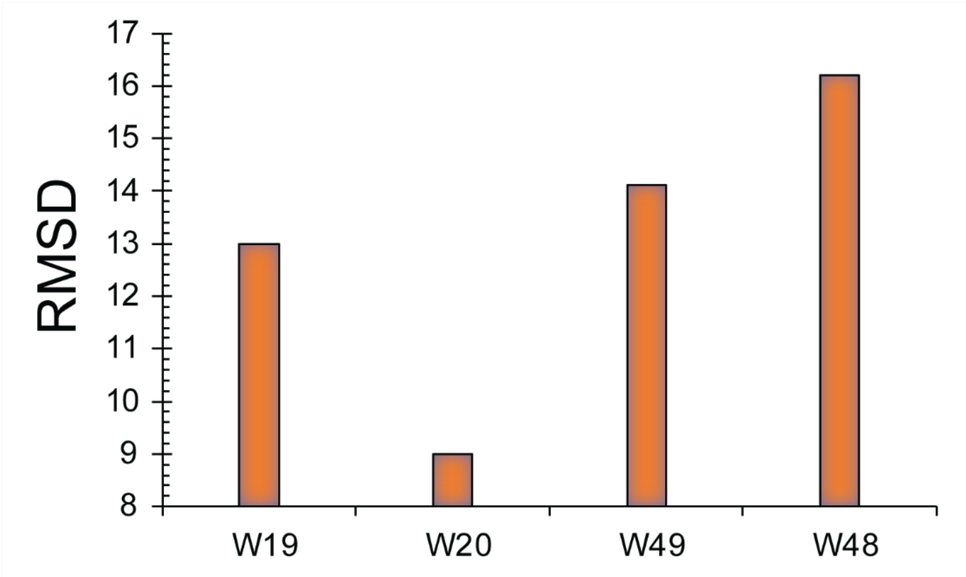

**Fig S8.The peak height of the waters at the beginning of the O4 channel from the OEC side.**

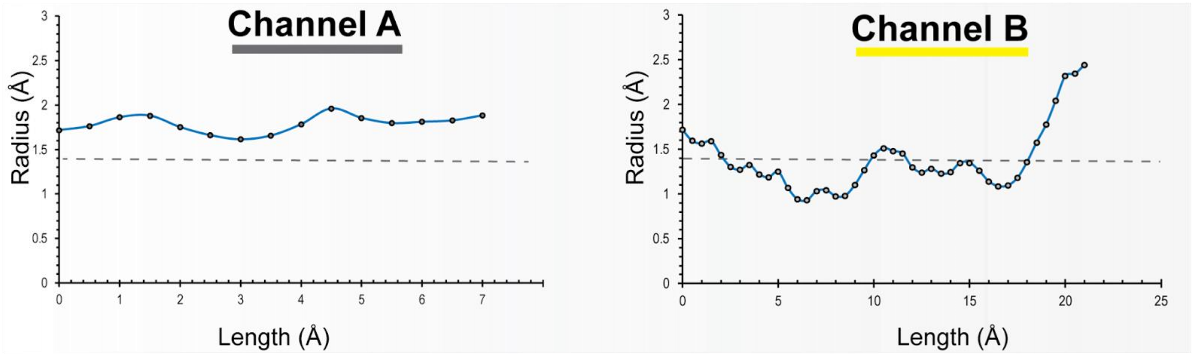

**Fig S9. Cavity radius of the channels connecting the BCT to the cytosol side.** The radius of the channels was calculated by CAVER. The gray dotted line represents the approximate radius of a water molecule.

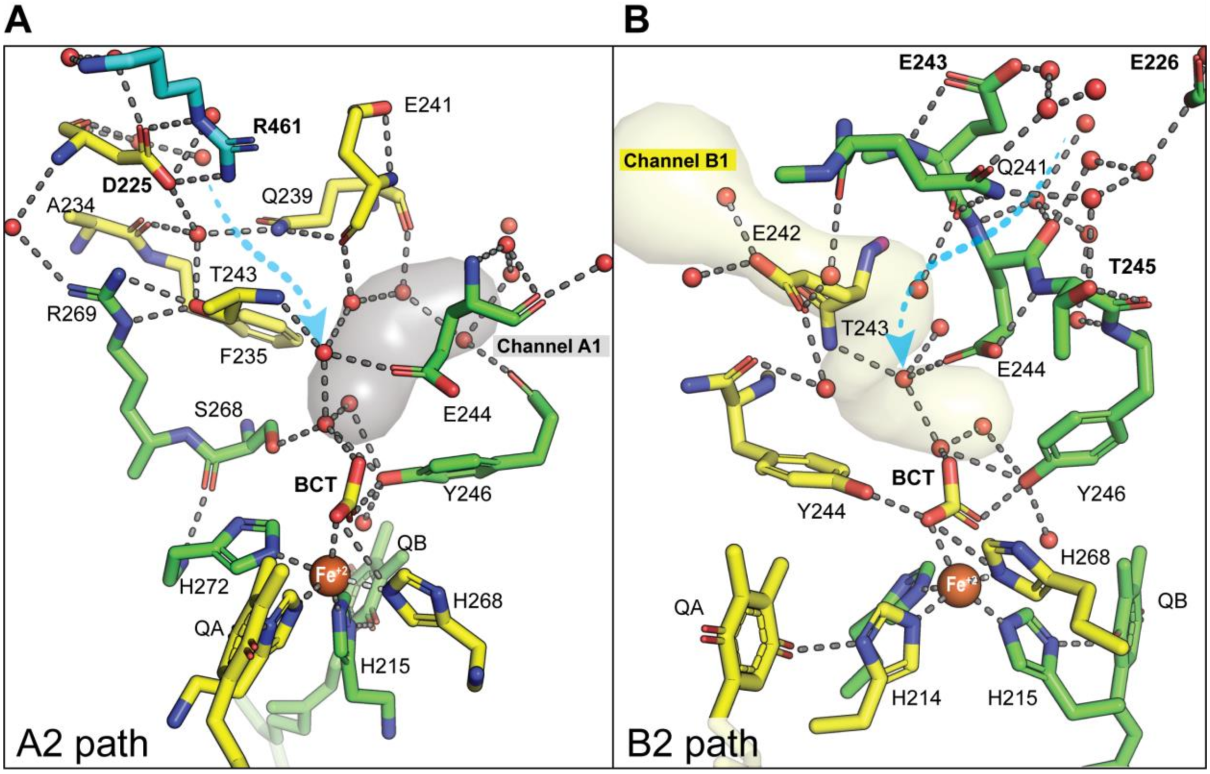

**Fig S10. Alternative Protonation Pathways for BCT.** The dotted blue arrows illustrate water channels proposed by a recent molecular dynamics (MD) simulation study (A2, B2) (*13*). However, based on our current structural analysis, these channels were not identified. Furthermore, the water structure in our model suggests that these channels are unlikely to serve as protonation pathways for the BCT. The interaction distance between atoms up to 3.2 Å is shown as a gray dashed line. Water channels generated by CAVER (channel A1 and channel B2) are colored as follows: channel A in gray and channel B in pale yellow.

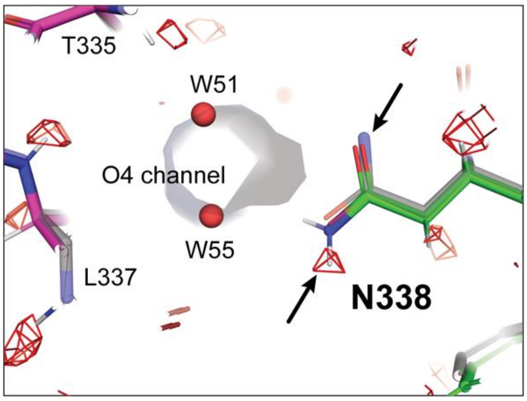

**Fig. S11 Impact of detecting hydrogen in modelling the amide group of D1-N338.** Hydrogen-omitted difference map at D1-N338 and the surrounding area. The omit map of the hydrogens and protons is shown as a red mesh and contoured at 8 σ. The black arrow highlights the nitrogen (N) position of the amide group in both the cryoEM structure (PDB ID: ...), and the RT XFEL structure (PDB ID: 7RF2) (*2*). The cryoEM structure is colored with residues present in D1 and CP43 subunits shown in green and magenta respectively, while the RT XFEL structure depicted in transparent gray in the background.

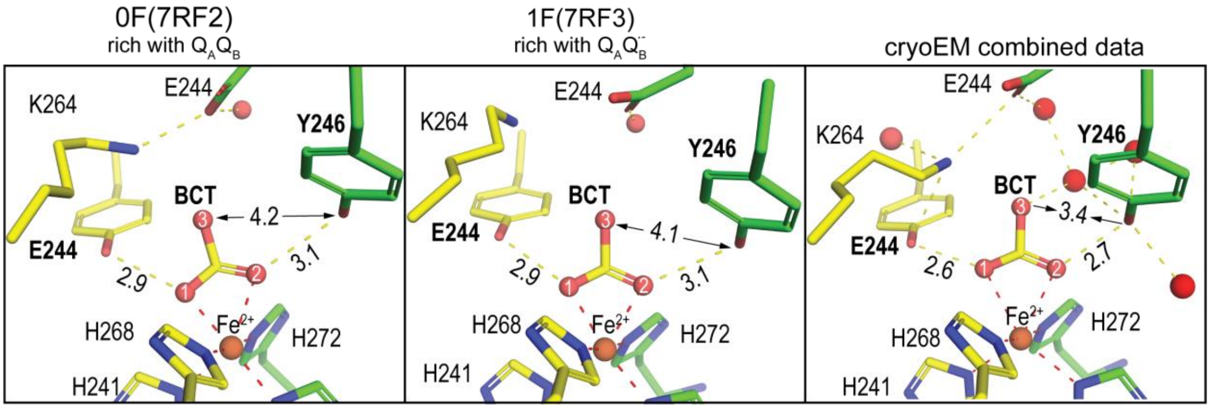

**Fig S12 The BCT and its surrounding environment in the current cryoEM model, with a comparative analysis of 0F and 1F structural data rich with $Q_AQ_B$ and $Q_AQ_B^-$ collected by RT XFEL (*2*).**

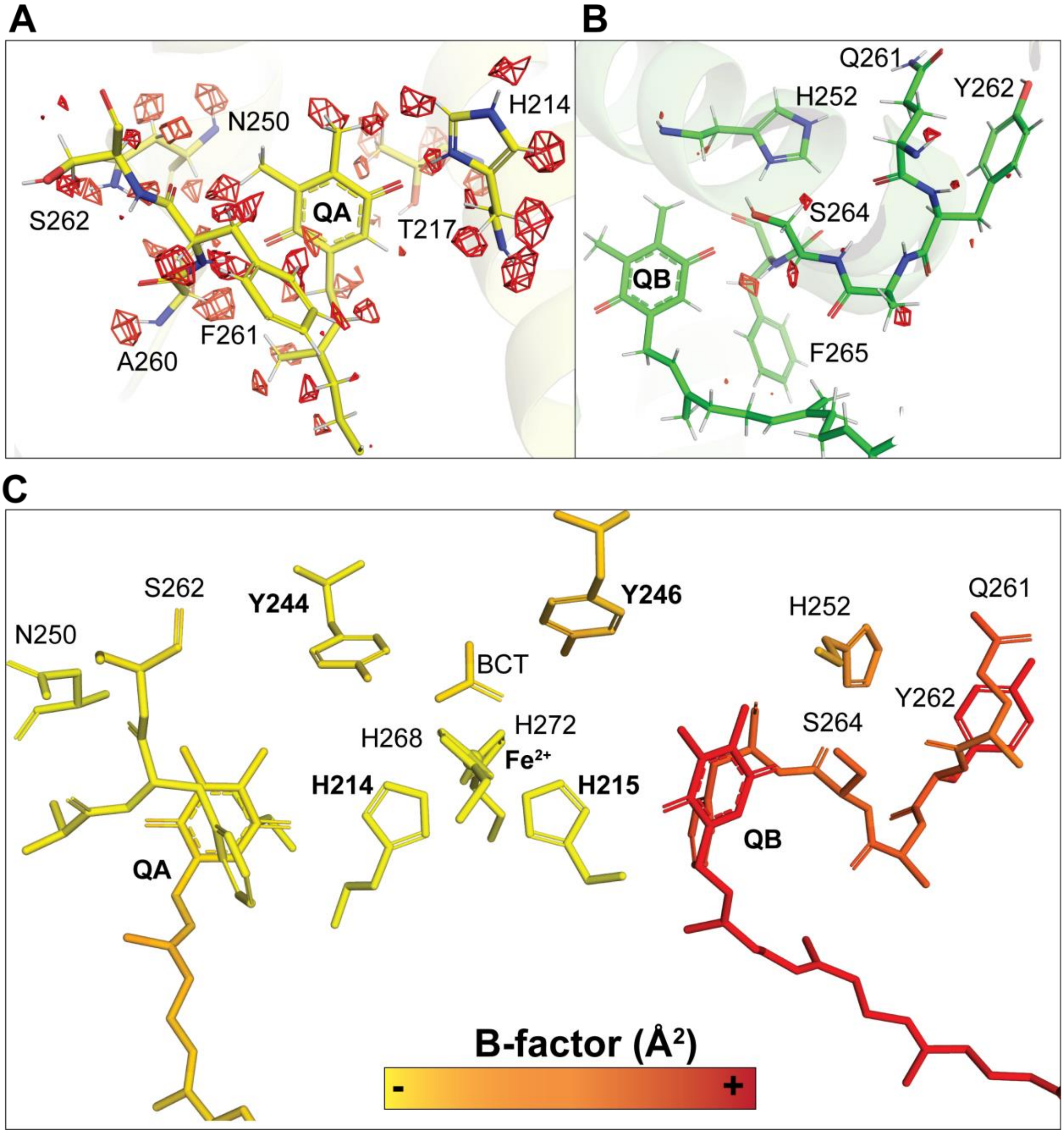

**Fig. S13. $Q_A$ and $Q_B$ sites. (A-B)** Omit difference map for hydrogen atoms at the $Q_A$ and $Q_B$ region. The omitted map of the hydrogens and protons is shown as a red mesh and contoured at 8 σ. **(C)** B-factor disruption at the $Q_A$ and $Q_B$ sites. The B-factor is color-coded, as described at the bottom middle, where 25 $Å^2$ is the lowest and 50 $Å^2$ is the highest value.

**Table. S3 Statistical analysis of detected Hydrogen peaks in the D1 subunit across different type bonds**

| Bond type[a] | Averged Distance (Å) | SD | Number of detected H-atoms | Averaged B-Factor ($Å^2$) of the parent atoms |
|---|---|---|---|---|
| **C-H[b]** | 1.19 | 0.25 | 539 | 27.8 |
| **$C-H_2$** | 1.25 | 0.26 | 519 | 28.8 |
| **$C-H_3$** | 1.37 | 0.26 | 437 | 27.7 |
| **N-H** | 1.18 | 0.24 | 332 | 27.6 |
| **O-H[c]** | 1.37 | 0.30 | 17 | 28.6 |

[a]Only for polymer in D1 subunit
[b]For alkyl and aromatic group
[c]Excluding side chain of glutamate and aspartate

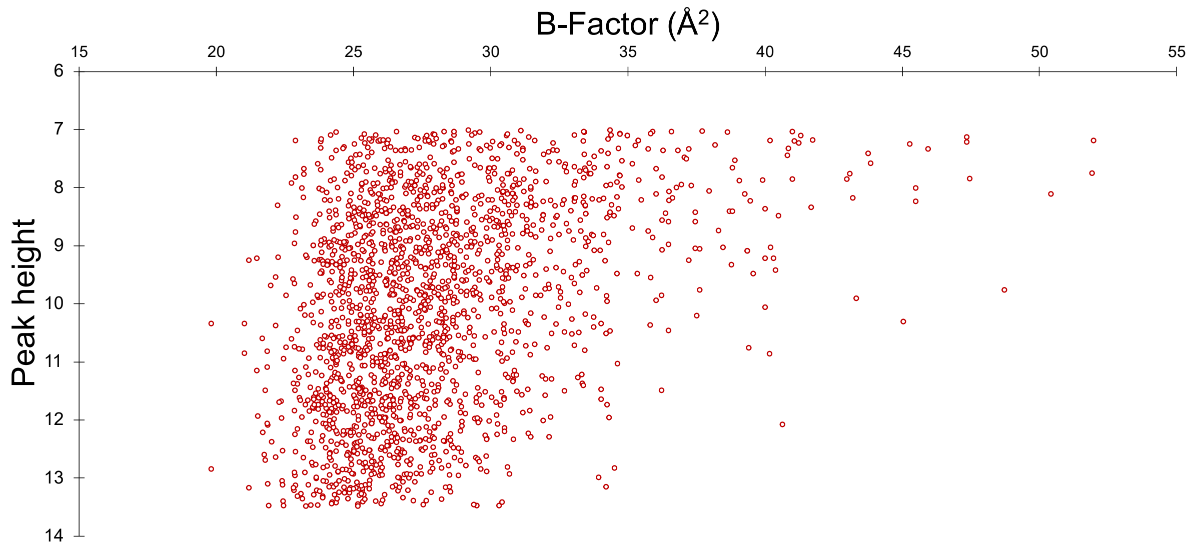

**Fig. S14. Chart shows the relation between the detected peak height of Hydrogens in the D1 subunit and the B-Factor of the parent atoms.**
